# Potent Antiviral Activities of Type I Interferons to SARS-CoV-2 Infection

**DOI:** 10.1101/2020.04.02.022764

**Authors:** Emily Mantlo, Natalya Bukreyeva, Junki Maruyama, Slobodan Paessler, Cheng Huang

## Abstract

The ongoing historic outbreak of COVID-19 not only constitutes a global public health crisis, but also carries a devastating social and economic impact. The disease is caused by a newly identified coronavirus, Severe Acute Respiratory Syndrome coronavirus 2 (SARS-CoV-2). There is an urgent need to identify antivirals to curtail the COVID-19 pandemic. Herein, we report the remarkable sensitivity of SARS-CoV-2 to recombinant human interferons α and β (IFNα/β). Treatment with IFN-α at a concentration of 50 international units (IU) per milliliter drastically reduces viral titers by 3.4 log or over 4 log, respectively, in Vero cells. The EC_50_ of IFN-α and IFN-β treatment is 1.35 IU/ml and 0.76 IU/ml, respectively, in Vero cells. These results suggest that SARS-CoV-2 is more sensitive than many other human pathogenic viruses, including SARS-CoV. Overall, our results demonstrate the potent efficacy of human Type I IFN in suppressing SARS-CoV-2 infection, a finding which could inform future treatment options for COVID-19.

## Introduction

The COVID-19 outbreak started in Wuhan, China in December 2019 and rapidly spread globally, causing over 752,000 confirmed cases and 36,000 deaths as of April 1, 2020. The causative agent for the COVID-19 disease is a newly identified Severe Acute Respiratory Syndrome coronavirus 2 (SARS-CoV-2) (1), which is transmitted through aerosol/droplet inhalation or contact. This historic outbreak has caused a public health crisis much more severe than the SARS outbreak, which “only” caused 8,098 infections and 774 deaths between November 2002 and July 2003. The COVID-19 outbreak also has had a devastating social and economic impact worldwide. In March, the World Health Organization has declared COVID-19 a pandemic. In the USA, there are over 200,000 confirmed cases and 4,300 deaths as of April 1, 2020. It is warned by the CDC that the COVID-19 pandemic may claim over 100,000 lives in USA (https://www.msn.com/en-nz/news/world/us-could-face-200000-coronavirus-deaths-millions-of-cases-fauci-warns/ar-BB11UlOj). Therefore, there is an urgent need to find treatments for COVID-19. Drugs already approved for the treatment of other diseases may offer the most expedient option for treating COVID-19, and several such drugs are already being tested in clinical trials.

Type I interferons (IFN-α/β) have broad spectrum antiviral activities against RNA viruses, which act by inducing an antiviral response across a wide range of cell types and mediating adaptive immune response. Humans produce 13 types of IFN-α and a singular IFN-β (2). Type I IFNs ultimately induces a number of interferon-stimulated genes (ISGs) which encode for a variety of antiviral effectors (3). Notably, IFN-β production leads to a positive feedback loop that further stimulates the expression of many of the IFN-α genes (4). Clinically, Type I IFNs have already been approved for use in the treatment of certain cancers, autoimmune disorders, and viral infections (hepatitis B and hepatitis C). We assessed the sensitivity of SARS-CoV-2 to both IFN-α and IFN-β *in vitro*. Herein, we report that type I IFNs exhibited potent anti-SARS-CoV-2 activities in cultured cells, demonstrating the therapeutic potency of type I IFNs for COVID-19.

## MATERIALS AND METHODS

### Virus and Cells

The SARS-CoV-2 (USA-WA1/2020) was obtained from The World Reference Center for Emerging Viruses and Arboviruses (WRCEVA), University of Texas Medical Branch, Galveston, TX. Stock virus was propagated by infecting Vero cells (ATCC CCL-81) at a low multiplicity of infection (MOI) of 0.0025. Three days after infection, supernatants were harvested and centrifuged at 2000 rpm for 5 min to remove cell debris. Stock virus was titrated with a 50% tissue culture infectious dose assay (TCID_50_) (5). All experiments involving infectious virus were conducted at the University of Texas Medical Branch (Galveston, TX) in approved biosafety level 3 laboratories in accordance with institutional health and safety guidelines and federal regulations.

### Virus growth curve

Vero cells were infected by SARS-CoV-2 at MOI 1 or 0.01 for 1 hr. Then inoculum was removed, replaced with media (DMEM+5%FBS) and incubated at 37 °C and 5% CO_2_. At different time points after infection, supernatants were harvested. Virus titers were determined by a TCID_50_ assay on Vero cells.

### Virus sensitivity to IFN treatment (infectious virus reduction assay)

Vero cells (2×10^4^/well) were seeded into 48-well plates for 24 h and treated with human IFN-β1a (mammalian, cat# 11415, PBL) and IFN-α (Universal Type I alpha A/D (Bg III), PBL, cat# 11200-1) at different concentrations for 16 h. Cells were then infected with SARS-CoV-2 at an MOI of 0.01 TCID_50_/cell. IFNs were supplemented after virus infection. Supernatants were collected at 22 hr post infection and assayed for virus titers.

### Virus sensitivity to IFN treatment (CPE inhibition assay)

Vero cells grown on 96-well plates (2×10^4^/well) were treated with 2-fold serial diluted human IFN-β1a or IFN-α for 16 h (250 IU/ml to 0.49 IU/ml). Cells were then infected with SARS-CoV-2 at an MOI of 0.01 TCID_50_/cell or Vesicular stomatitis virus (VSV, Indiana strain) at MOI 0.1 PFU/cell for 1 hr. The inoculums were removed and replaced with fresh media. As controls, cells were mock-infected, or infected without IFN treatment. All experiments were performed in quadruplicates. For VSV samples, the supernatants were aspirated at 12 hpi. The monolayers were washed with PBS for three times to remove dead cells, fixed with 10% formaldehyde, and stained with crystal violet for cytopathic effect (CPE) observation. For SARS-CoV-2 samples, CPE was observed at 72 hpi.

## Results

The growth kinetics of the newly identified SARS-CoV-2 in cultured cells had not been characterized. Thus, we first examined the growth kinetics of SARS-CoV-2 in Vero cells. Vero cells were infected at either a low MOI (MOI=0.01) or high MOI (MOI=1). Supernatant was collected every 8-16 hours. At both conditions, viral titers peaked at approximately 24 hours post-infection (hpi) and remained stable until 40 hours post-infection before declining (Fig. 1). The peak virus titer was 5.5×10^6^ TCID_50_/ml at MOI 0.01 and 3.75×10^5^ TCID_50_/ml at MOI 1, indicating that viral replication was more efficient at a low MOI (MOI=0.01) than a high MOI (MOI=1). Additionally, virus infection caused strong cytopathic effect (CPE), which was evident at 48 hpi, much later than the peak of virus production (at 40 hpi).

**Figure 1:**
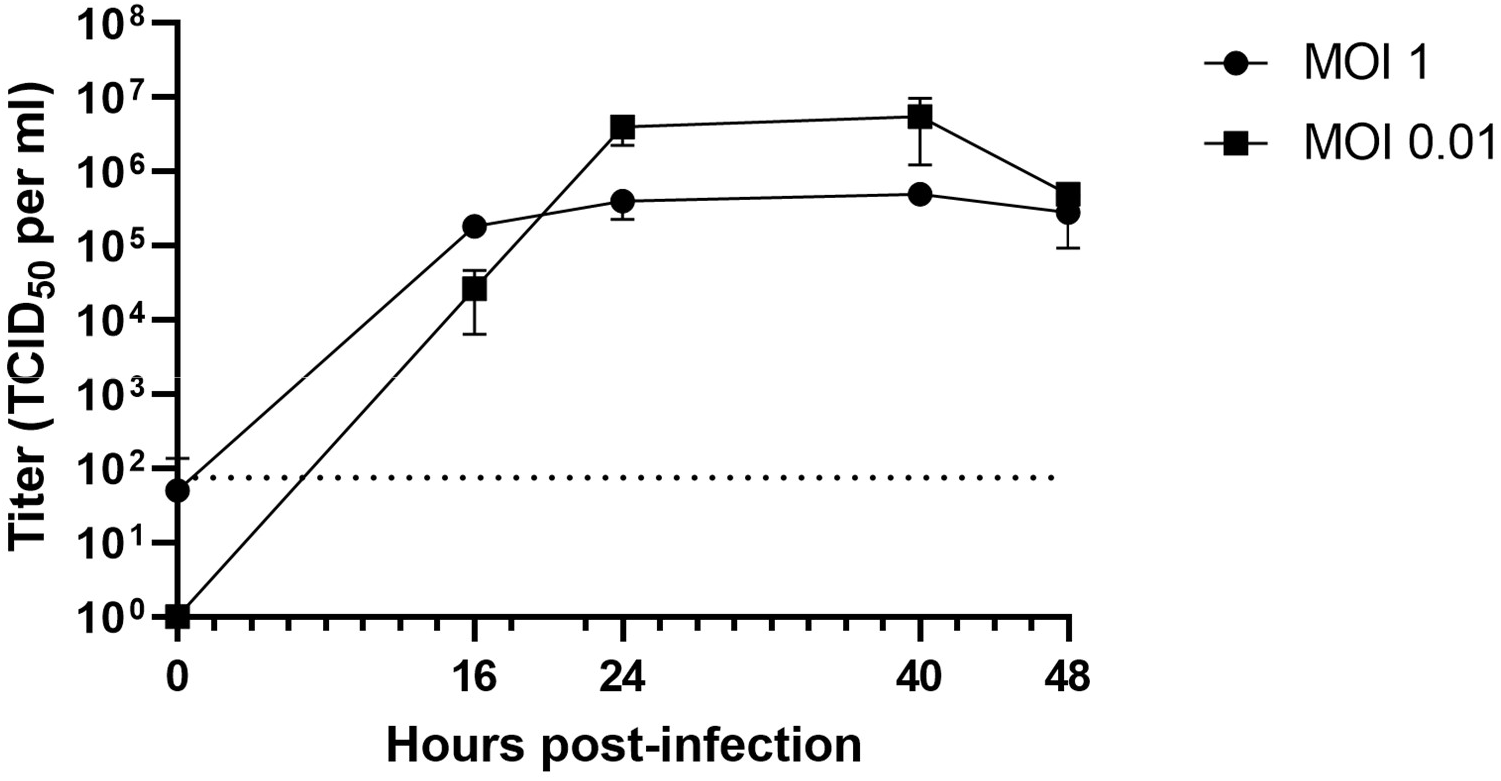
Vero cells were infected by SARS-CoV-2 at MOI 1 or 0.01 for 1 hr. At different time points after infection, virus titers were determined by a TCID_50_ assay on Vero cells. The average of triplicates and Standard deviation are shown. Dotted line indicates the detection limit.

Next, we examined the effect of recombinant human IFN-α and IFN-β treatment on viral infection. Vero cells were pre-treated with different concentrations of IFN-α or IFN-β ranging from 50-1000 international units (IU) per milliliter for 16 hours. After 1 hour of infection with SARS-CoV-2 (MOI 0.01), media containing IFN was returned, and cells were incubated for a further 22 hours. Supernatants were then collected, and viral titers were determined via TCID_50_ assay. The result indicated that IFN-α treatment potently inhibited SARS-CoV-2 infection. Virus titers were not detectable except at the lowest concentration tested (50 IU/ml), at which the viral titers were drastically reduced by 4 logs of magnitude (Fig. 2). For IFN-β, the virus titers were below the detection limit at all concentrations tested (50 u/ml-1000u/ml), indicating more potent anti-SARS-CoV-2 activity than IFN-α. Consistently, no CPE was observable under microscopic examination in all IFN-treated samples.

**Figure 2:**
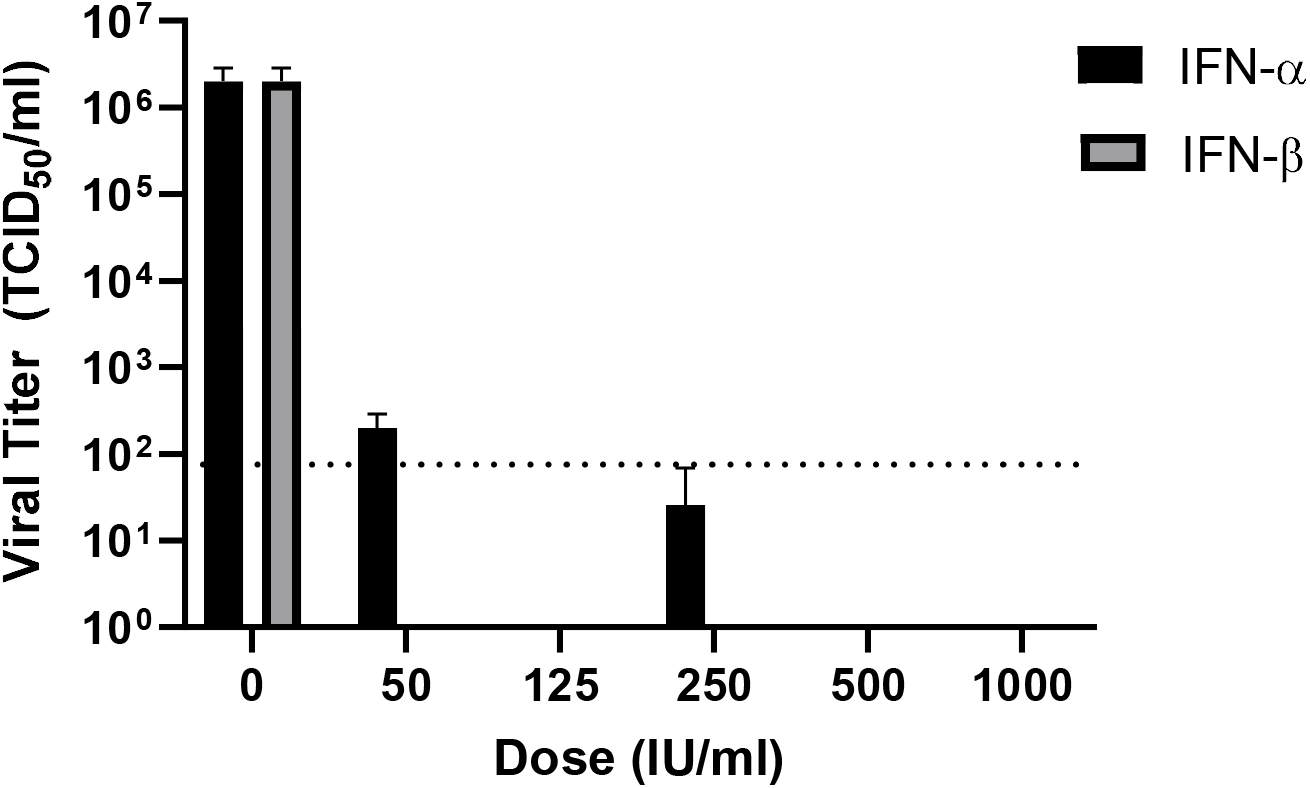
Vero cells were pretreated with human IFN-α or IFN-β (0, 50, 125, 250, 500, 1000 IU/ml) for 16 hours, and then infected with SARS-CoV2 for 1 hour at an MOI of 0.01. Viral inoculums were removed and replaced with fresh media containing listed concentrations of IFN-α or IFN-β. Media was collected at 22 hpi and titers were determined via TCID50 assay on Vero cells. The average of triplicates and Standard deviation are shown. Dotted line indicates the detection limit.

We next tested the antiviral efficacy of IFN-α and IFN-β at lower concentrations (1-50 IU/ml). Both IFN-α and IFN-β dose-dependently inhibited virus infection at these lower concentrations (Fig. 3). IFN-α exhibited anti-SARS-CoV-2 activity at a concentration as low as 5 IU/ml, resulting in a significant reduction of viral titer by over 1 log (P<0.01). With increasing IFN-α concentrations, the virus titers steadily decreased. Treatment with IFN-α at 50 IU/ml drastically reduces viral titers by 3.4 log. Treatment with 1 IU/ml of IFN-β resulted in a moderate (approximately 70%) but significant decrease in virus titer (P<0.05, Student t test). Infectious virus was nearly undetectable upon treatment with 10, 25, and 50 IU/ml of IFN-β. The EC_50_ of IFN-α and IFN-β treatment is 1.35 IU/ml and 0.76 IU/ml, respectively. Taken together, these results indicate that treatment with low concentrations of both IFN-α and IFN-β significantly inhibited viral infection, with IFN-β being slightly more effective than IFN-α.

**Figure 3:**
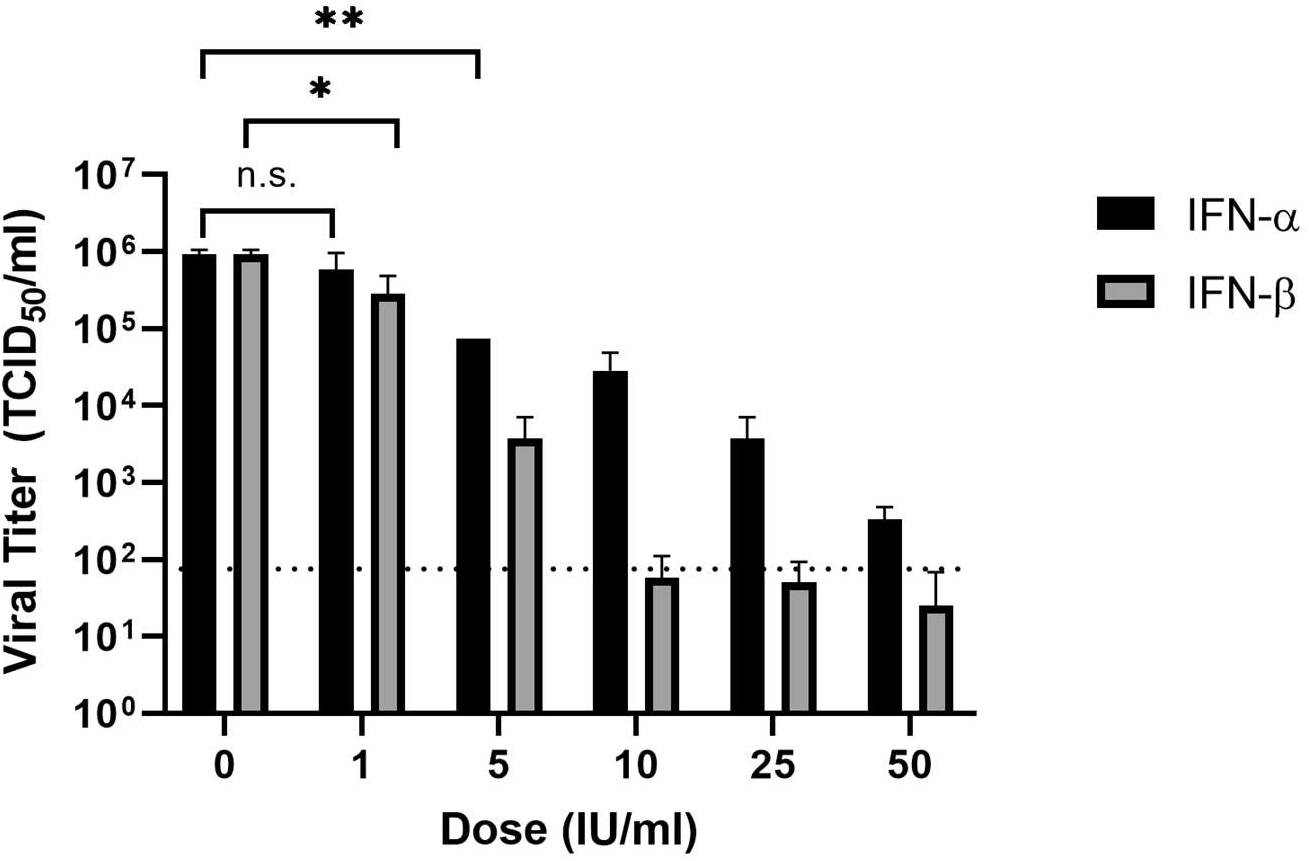
Vero cells were pretreated with human IFN-α or IFN-β (0, 1, 5, 10, 25, 50 U/ml) for 16 hours and then infected with SARS-CoV-2 at an MOI of 0.01. Viral inoculums were removed and replaced with fresh media containing listed concentrations of IFN-α or IFN-β. Virus titers at 22 hpi were determined via TCID50 assay. The average of triplicates and Standard deviation are shown. Dotted line indicates the detection limit. (*, P<0.05; **, P<0.01; n.s. not significant, one tail Student T test)

In addition, we compared the IFN sensitivity of SARS-CoV-2 with that of Vesicular stomatitis virus (VSV), an IFN-sensitive RNA virus. IFN-α or IFN-β were 2-fold serially diluted (250 IU/ml to 0.49 IU/ml) and added to Vero cells for 16 hr. Then cells were infected by VSV (MOI 0.1) or SARS-CoV-2 (MOI 0.01). CPE were observed at 12 hpi for VSV and 72 hpi for SARS-CoV-2. In VSV-infected cells, IFN-α and IFN-β both inhibited CPE development at a concentration of 31.25 IU/ml, while at 15.6 IU/ml the CPE was not discernable from that of IFN-untreated samples. For SARS-CoV2, the lowest concentration that IFN-β or IFN-α inhibited CPE was 31.25 IU/ml and 62.5 IU/ml, respectively. The CPE inhibition data suggests that the IFN sensitivity of SARS-CoV-2 is comparable to that of VSV.

## Discussion

Our data clearly demonstrate that SARS-CoV-2 is highly sensitive to both IFN-α and IFN-β treatment in cultured cells, which is comparable to the IFN-sensitive VSV. Our discovery reveals a weakness of the new coronavirus, which may be informative to antiviral development. The experiment was performed in the IFN-α/β gene-defective Vero cells (6). It is plausible that in IFN-competent cells the efficacy of exogenous IFN-β treatment against SARS-CoV-2 infection is more potent, as IFN-β upregulates other subtypes of Type I IFN expression and augments the IFN-mediated antiviral response (4). Our data may provide an explanation, at least in part, to the observation that approximately 80% of patients actually develop mild symptoms and recover (7). It is possible that many of them are able to mount IFN-α/β-mediated innate immune response upon SARS-CoV-2 infection, which helps to limit virus infection/dissemination at an early stage of disease. At a later stage, the adaptive immune response (antibody etc.) may eventually help patients recover from the COVID-19 disease.

Compared to SARS-CoV-2, it seems that SARS-CoV is relatively less sensitive to IFN treatment *in vitro* (8, 9). One study reported that the EC_50_ of IFN-β for SARS-CoV is 95 or 105 IU/ml depending on virus strains (10). Many other highly pathogenic viruses are also resistant to exogenous IFN treatment. For Ebola virus, it has been reported that treatment with exogenous IFN-α does not affect viral replication and infectious virus production in cultured cells (11), probably as a result of antagonism of the IFN response by viral protein. Junín virus, an arenavirus that causes Argentine Hemorrhagic Fever, is likewise insensitive to IFN treatment. When treated with a high concentration of human IFN-α, β or γ (1000 U/ml), the titers of JUNV were reduced by less than 1-log in Vero cells. Further work is warranted to characterize the IFN response during SARS-CoV-2 infection to better understand the underlying mechanism behind its IFN sensitivity.

*In vitro*, we have demonstrated that SARS-CoV-2 replication is inhibited by IFN-α and IFN-β at concentrations that are clinically achievable in patients. Recombinant IFN-αs, Roferon-A and Intron-A, which have been approved for hepatitis B and C treatment, can reach concentrations of up to 330 IU/ml and 204 IU/ml, respectively, in serum (12). Recombinant IFN-β drugs, Betaferon and Rebif, which have been approved for the treatment of multiple sclerosis, can reach concentrations of 40 IU/ml and 4.1 IU/ml, respectively, in serum (12). Therefore, some of these drugs may have the potential to be repurposed for the treatment of COVID-19 either alone or in combination with other antiviral therapies.

## Acknowledgments

We thank Drs. Kenneth Plante (The World Reference Center for Emerging Viruses and Arboviruses, UTMB) and Natalie Thornburg from the CDC for providing the SARS-CoV-2 stock virus. Work in the Paessler laboratory was supported in parts by Public Health Service grants R01AI093445 and R01AI129198 and the John. S. Dunn Distinguished Chair in Biodefense endowment. C.H. was supported by UTMB Commitment Fund P84373 and Department of Pathology Pilot Grant and would like to acknowledge the Galveston National Laboratory (supported by the Public Health Service award 5UC7AI094660) for support of his research activity.

## References

1. Zhou P, Yang XL, Wang XG, Hu B, Zhang L, Zhang W, Si HR, Zhu Y, Li B, Huang CL, Chen HD, Chen J, Luo Y, Guo H, Jiang RD, Liu MQ, Chen Y, Shen XR, Wang X, Zheng XS, Zhao K, Chen QJ, Deng F, Liu LL, Yan B, Zhan FX, Wang YY, Xiao GF, Shi ZL. A pneumonia outbreak associated with a new coronavirus of probable bat origin. Nature. 2020;579(7798):270–3. Epub 2020/02/06. doi: 10.1038/s41586-020-2012-7. PubMed PMID: 32015507; PMCID: PMC7095418.

2. Pestka S, Krause CD, Walter MR. Interferons, interferon-like cytokines, and their receptors. Immunol Rev. 2004;202:8–32. Epub 2004/11/18. doi: 10.1111/j.0105-2896.2004.00204.x. PubMed PMID: 15546383.

3. Schoggins JW, Wilson SJ, Panis M, Murphy MY, Jones CT, Bieniasz P, Rice CM. A diverse range of gene products are effectors of the type I interferon antiviral response. Nature. 2011;472(7344):481–5. Epub 2011/04/12. doi: 10.1038/nature09907. PubMed PMID: 21478870; PMCID: PMC3409588.

4. Honda K, Takaoka A, Taniguchi T. Type I interferon [corrected] gene induction by the interferon regulatory factor family of transcription factors. Immunity. 2006;25(3):349–60. Epub 2006/09/19. doi: 10.1016/j.immuni.2006.08.009. PubMed PMID: 16979567.

5. Narayanan K, Huang C, Lokugamage K, Kamitani W, Ikegami T, Tseng CT, Makino S. Severe acute respiratory syndrome coronavirus nsp1 suppresses host gene expression, including that of type I interferon, in infected cells. J Virol. 2008;82(9):4471–9. Epub 2008/02/29. doi: 10.1128/JVI.02472-07. PubMed PMID: 18305050; PMCID: PMC2293030.

6. Desmyter J, Melnick JL, Rawls WE. Defectiveness of interferon production and of rubella virus interference in a line of African green monkey kidney cells (Vero). J Virol. 1968;2(10):955–61. Epub 1968/10/01. PubMed PMID: 4302013; PMCID: PMC375423.

7. Wu Z, McGoogan JM. Characteristics of and Important Lessons From the Coronavirus Disease 2019 (COVID-19) Outbreak in China: Summary of a Report of 72314 Cases From the Chinese Center for Disease Control and Prevention. JAMA. 2020. Epub 2020/02/25. doi: 10.1001/jama.2020.2648. PubMed PMID: 32091533.

8. Moriguchi H, Sato C. Treatment of SARS with human interferons. Lancet. 2003;362(9390):1159. Epub 2003/10/11. doi: 10.1016/S0140-6736(03)14484-4. PubMed PMID: 14550718.

9. Dahl H, Linde A, Strannegard O. In vitro inhibition of SARS virus replication by human interferons. Scand J Infect Dis. 2004;36(11-12):829–31. Epub 2005/03/15. doi: 10.1080/00365540410021144. PubMed PMID: 15764169.

10. Cinatl J, Morgenstern B, Bauer G, Chandra P, Rabenau H, Doerr HW. Glycyrrhizin, an active component of liquorice roots, and replication of SARS-associated coronavirus. Lancet. 2003;361(9374):2045–6. Epub 2003/06/20. doi: 10.1016/s0140-6736(03)13615-x. PubMed PMID: 12814717.

11. Kash JC, Muhlberger E, Carter V, Grosch M, Perwitasari O, Proll SC, Thomas MJ, Weber F, Klenk HD, Katze MG. Global suppression of the host antiviral response by Ebola- and Marburgviruses: increased antagonism of the type I interferon response is associated with enhanced virulence. J Virol. 2006;80(6):3009–20. Epub 2006/02/28. doi: 10.1128/JVI.80.6.3009-3020.2006. PubMed PMID: 16501110; PMCID: PMC1395418.

12. Strayer DR, Dickey R, Carter WA. Sensitivity of SARS/MERS CoV to interferons and other drugs based on achievable serum concentrations in humans. Infect Disord Drug Targets. 2014;14(1):37–43. Epub 2014/07/16. doi: 10.2174/1871526514666140713152858. PubMed PMID: 25019238.

